# A novel cell segmentation method for developing embryos using machine learning

**DOI:** 10.1101/288720

**Authors:** Rikifumi Ota, Takahiro Ide, Tatsuo Michiue

## Abstract

Cell segmentation is crucial in the study of morphogenesis in developing embryos, but it is limited in its accuracy. In this study we provide a novel method for cell segmentation using machine-learning, termed Cell Segmenter using Machine Learning (CSML). CSML performed better than state-of-the-art methods, such as RACE and watershed, in the segmentation of ectodermal cells in the *Xenopus* embryo. CSML required only one whole embryo image for training a Fully Convolutional Network classifier, and it took 20 seconds per each image to return a segmented image. To validate its accuracy, we compared it to other methods in assessing several indicators of cell shape. We also examined the generality by measuring its performance in segmenting independent images. Our data demonstrates the superiority of CSML, and we expect this application to significantly improve efficiency in cell shape studies.

## Introduction

In multicellular organisms, groups of cells cooperatively behave and change morphology, resulting in dynamic tissue deformation. In vertebrate gastrula, for example, coordinated migration of mesodermal cells occurs, including convergent extension, to establish complex structures inside the embryo to complete gastrulation (Keller and Danilchik, 1988). To understand tissue deformation, it is necessary to describe the overall cell arrangement. Several methods for describing cell arrangement have been reported (Merkel and Manning, 2016) that use strategies such as the relative position of the cell center (Guirao et al., 2015), the vertical cell shape by fitting of an ellipse (Blanchard et al., 2009), or direct tracing of the cell shape via watershed cell segmentation, which provides the most information, including the apical cell area.

Watershed cell segmentation is one of the most commonly used tools to segment touching cells (Meijering, 2012), where the image is segmented into catchment basins and watershed ridge lines based upon light and dark pixels (Beucher, 1982). Real-time Accurate Cell-shape Extractor (RACE) is a state-of-the-art method for three-dimensional cell segmentation using slice-based segmentation approaches (Stegmaier et al., 2016). Seed-based watershed for cell delineation has been developed with pattern-recognition methods to track dividing cells (Wang et al., 2017).

The motivation for utilizing these methods arises from the difficulty in describing the actual cell shape. Cell shape data is usually obtained by labeling the cell membrane with either membrane-bound GFP in living cells or phalloidin staining in fixed cells, and then the fluorescent image is transformed into a line drawing for subsequent analyses. Ordinary watershed methods often produce over-segmentation that require further processing (Meijering, 2012). To overcome this inconvenience, we applied machine learning to the watershed method, specifically, a Fully Convolutional Network (FCN) that efficiently learns semantic segmentation from spatial output maps (Long, 2015).

In this study we used machine learning for cell segmentation to achieve high quality segmentation. The FCN classifier was used to classify each pixel by semantic segmentation into one of two classes: membrane or non-membrane. Our approach provided the highest accuracy of the three methods compared: Cell Segmenter using Machine Learning (CSML), RACE, and watershed. Cell shape data using CSML was close to Ground Truth. Our approach was also validated for generality, and the program took 20 seconds to return each segmented image. As such, our CSML program will greatly improve the efficiency of obtaining high-quality cell segmentation data.

## Materials and Methods

### Collection of cell shape images in *Xenopus* embryos

*Xenopus laevis* embryos were used to obtain cell boundary contoured images, where 0.5 ng of membrane GFP (mGFP) mRNA was injected into the animal region of four-cell embryos. mGFP mRNA was transcribed *in vitro* with mMessage mMachine SP6 kit using mGFP-CS2 as a template. Injected embryos were developed until mid-gastrula stage, and then fluorescent images were taken by confocal laser microscopy (Olympus, Tokyo). With this approach, fluorescent images of ectodermal cells were obtained in most cases.

### Calculation of cell shape

Area, perimeter, solidity, eccentricity and orientation (inferred by regionprops, MATLAB) were used as indicated, with all data collected using MATLAB. Solidity is the proportion of pixels in the smallest convex polygon that can contain the region. Eccentricity is ratio of the distance between the major axis length and the foci of the ellipse that has the same second-moments as the region. Orientation is the angle between the x-axis and the major axis of the ellipse described in eccentricity.

## Results

### Design of a novel cell segmentation application

Our cell segmentation application using machine learning was firstly developed. Then, to validate the quality of segmentation by CSML, we also created a watershed transform program. CSML consists of three phases (Fig. 1). First, small images are obtained by preprocessing a whole image. Then, inferred small images are obtained using an FCN classifier. Lastly, a whole segmentation image is obtained by postprocessing the inferred small images. Preprocessing before inference of images by the FCN classifier enables all non-overlapping small (s x s pixels, s=32 in this application), square images cut out from the whole image to be normalized by succession of standardization, sigmoid filter and standardization to enhance membranes. In postprocessing, after acquiring binary small images, one inferred whole image is obtained by merging two inferred images. For the merge, we adopted the maximum value of two inferred images. One is an inferred whole image made by the connection of s x s pixel binary images, and the other is made by the connection of s/2-pixel-shifted s x s pixel binary images. The segmentation image is acquired by discarding the isolated area under 1000 px inferred as membrane, dilating the inferred image by one pixel, filling the area under 1 px to remove noise, and watershedding. This overlapping process is essential, as inferred s x s pixel images exhibit lower accuracy in the periphery.

**Figure 1.**
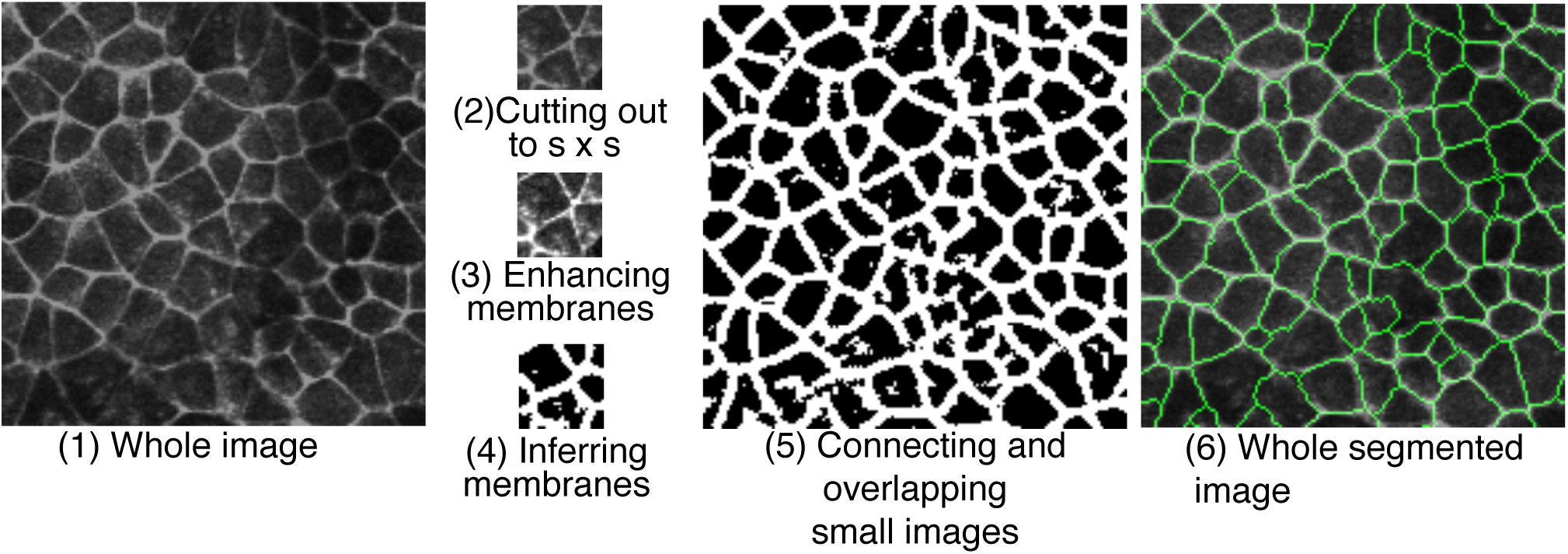
Outline of CSML segmentation procedure to acquire an inferred image. Binary small images were obtained by taking s x s small images from the whole image. Membranes were enhanced, and then the images were classified with the FCN classifier. A whole segmented image was obtained by overlapping two whole images, one being connected small images and the other s/2-pixel-shifted small images, and then removing noise and watershedding. In the training step, images were processed in the same way using enhanced small images.

The FCN classifier computes the probability of each pixel in the input image being in the membrane class. There are two steps in the algorithm: a training step and an inference step. In the training step, training images with corresponding manually segmented images are provided to the FCN classifier as input, and the weights of the FCN classifier are optimized. For training, 24,500 images of s x s images taken from one whole gray-scale training image were used. Training images were picked from 24 regions of randomly selected 2s × 2s sizes that spanned the whole image. In the inference step, only images are given to the FCN classifier and inferred images are returned (Fig. 1). The FCN classifier consists of 13 layers in the following order: two convolutional and relu layers, one batch normalization layer, one max pooling layer, two convolutional and relu layers, one batch normalization layer, one convolutional layer to reduce channels, and one deconvolutional layer. The convolutional layer outputs a map of dots convolving around an input map with multiple learned filters. The max pooling layer outputs a map of dots where each is a maximum of local non-overlapping square maps. The convolutional layer to reduce channels outputs one layer from multiple layers of input. The deconvolutional layer outputs a map by densifying the sparse map of input with multiple learned filters. The relu layer outputs same size of map of dots, where each has a function max (0, x) adapted to it.

In the watershed transform program, all non-overlapping s × s pixel square images were cut out from the whole image, and each image was binarized by the Otsu method (Otsu, 1979). The segmentation image was obtained by dilating the connected whole image by one pixel, filling the area under 10 px, and watershedding.

### Validation of the novel, machine learning-based program for cell segmentation

In order to assess the performance of this program, we compared segmentation images produced by CSML, RACE and watershed with manually annotated segmentation (Ground Truth), and measured the total time required to train the network and infer images in CSML and process the watershed transform program. In this validation, images used in the training step and the inference step were from the same embryo but at different developmental stages. We used segmentation images produced by RACE in the first step as 2D cell segments. We computed the accuracy of each method using the evaluation methods recommended by Coelho et al. (2009) (Fig. 2A). Precision and recall was calculated as 1-false positive rate and 1-false negative rate, respectively. The F-value was calculated as 2*precision*recall/(precision+recall). Higher precision indicates fewer split cells, whereas higher recall indicates fewer added cells (Coelho et al., 2009). We found that CSML provided better performance than RACE and watershed, as indicated by a higher F-value (Fig. 2A). Precision, recall, and F-value of CSML was 96%, 95% and 95%, respectively. The false-positive and false-negative error rates of CSML were only 4% and 5%, respectively. These comparisons show that CSML provided the highest recall and watershed provided the highest precision.

**Figure 2.**
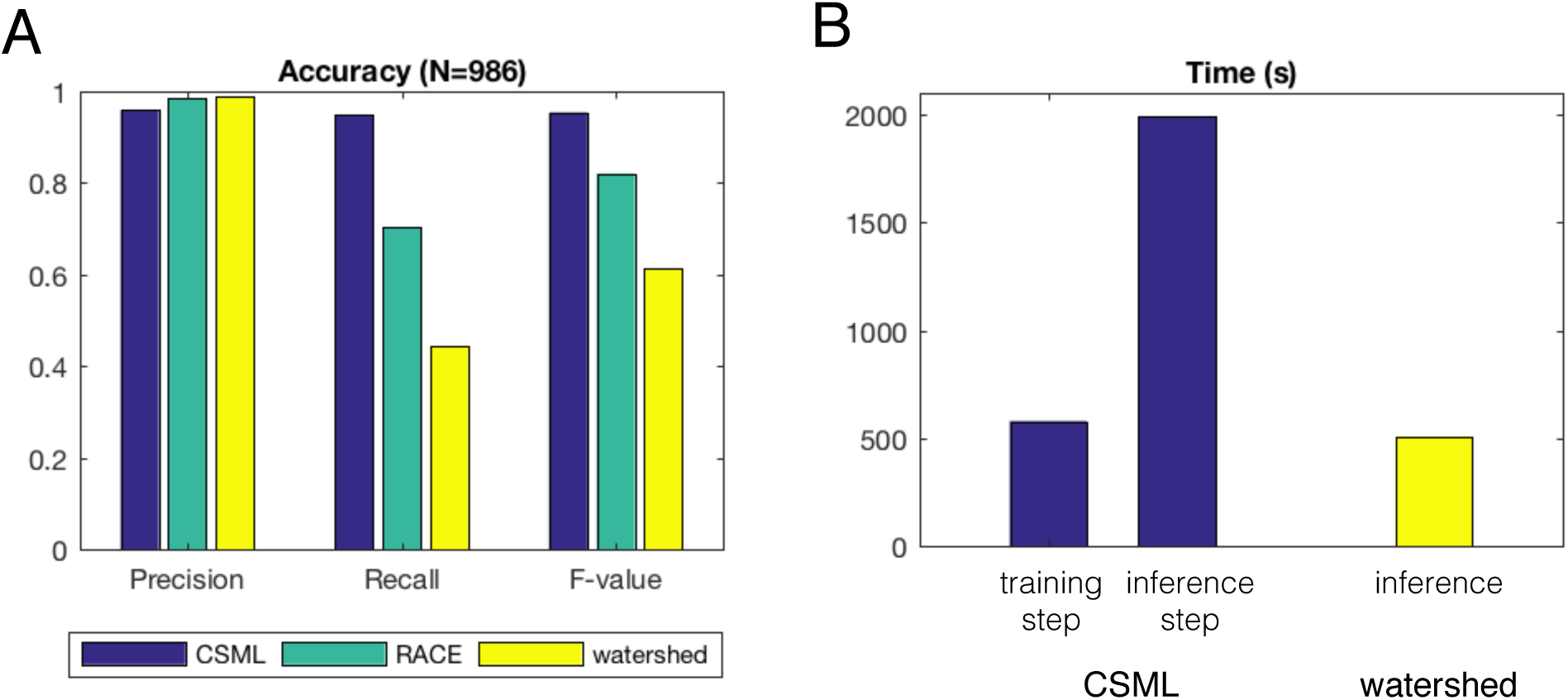
Assessment of accuracy and speed of CSML, RACE and watershed. (A) Comparison of precision, recall, and F-value of segmentation by CSML, RACE and watershed. CSML provided a higher F-value (better performance) than RACE and watershed. (B) Comparison of time required for segmentation. The performance was measured on an Apple MacBook Pro equipped with an 2.7 GHz Intel Core i5, 8GB memory and MacOS High Sierra 10.13.

To train the network, it took around 600 seconds using 24,500 small training images, which is required only once before the inference. To infer 100 images (1024 px × 1024 px), CSML took less than 2,000 seconds, whereas the watershed transform program required around 500 seconds (Fig. 2B).

### Validation of cell shape data

Due to the high quality of segmentation, cell shape data can be directly extracted from segmented images with CSML. In order to assess the accuracy in cell shape data, several features of the shape of cells in the segmentation images produced by CSML, RACE, watershed were compared with those produced by Ground Truth. In area and perimeter, CSML showed the best consistency to Ground Truth (Fig. 3A, B), whereas RACE and watershed generally showed a greater distribution of cells with large values. Furthermore, cells with very small values were observed in the watershed-derived data. This could be because both RACE and watershed had a tendency of failing to recognize cells, and watershed further performs excessive segmentation to describe very small sized cells. Cell solidity by RACE and watershed possessed lower values than CSML and Ground Truth (Fig. 3C). Again, the former failed to recognize some cells, regarded several cells as one, and found cells to be more dented. In eccentricity, CSML and RACE showed the best correspondence to Ground Truth (Fig. 3D), while the graph of watershed was shifted to right due to a tendency to merge cells to be thin. In orientation, CSML showed the best correspondence to Ground Truth, but it was not particularly close (Fig. 3E). RACE showed a little similarity with Ground Truth, and the graph of watershed was obviously different. Taken together, these results show that CSML had the best performance among the three cell segmentation methods in relation to cell shape features.

**Figure 3.**
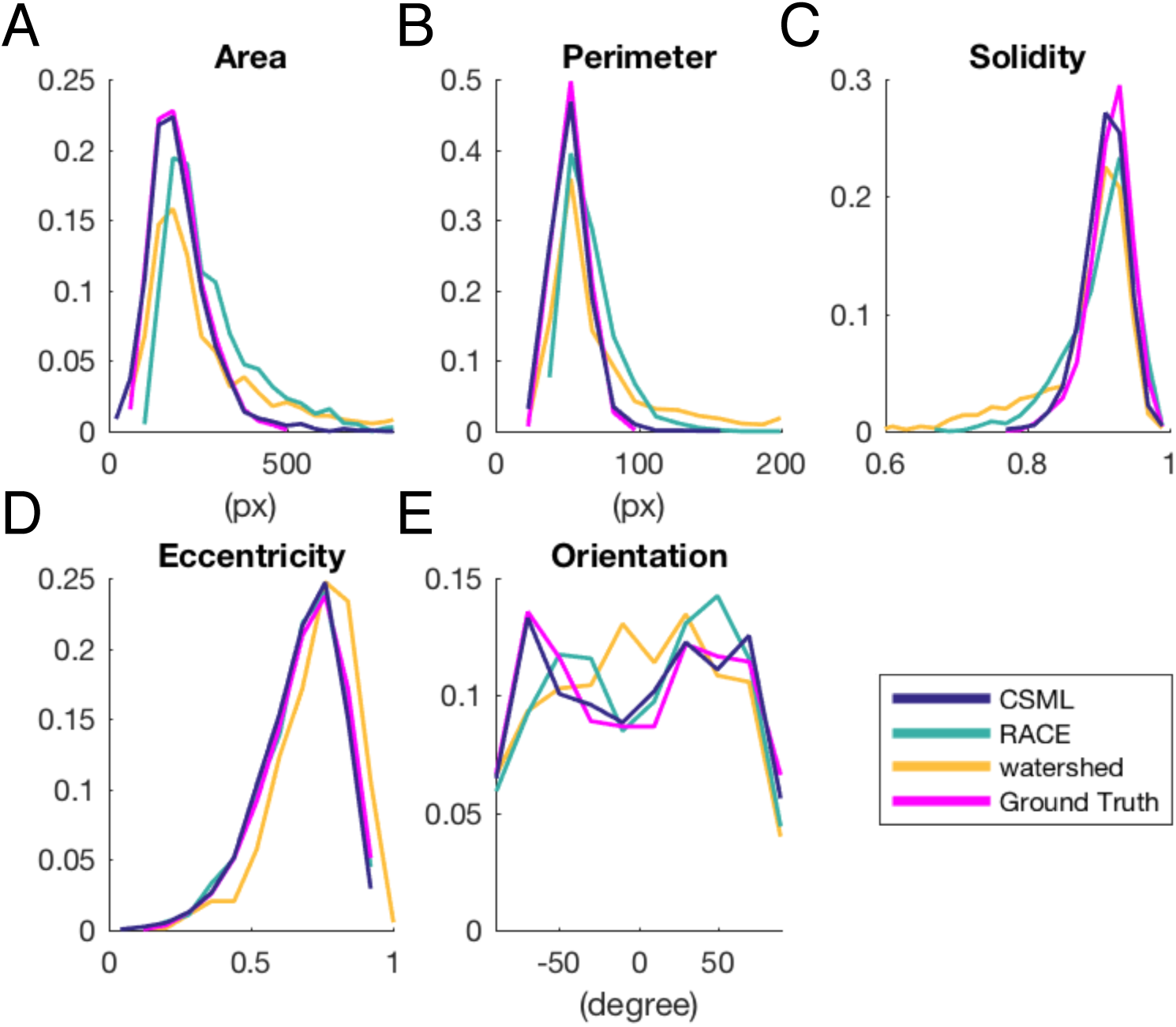
Comparison of cell shape features in segmentation images produced by CSML, RACE, and watershed with Ground Truth. Area (A), perimeter (B), solidity (C), eccentricity (D) and orientation (E) were used as indicators. The graph of area and perimeter for RACE and watershed had a shift to the right.

### Influence of changed training conditions on accuracy

In order to observe how the accuracy was affected by training conditions, we compared segmentation images produced from various numbers and types of training images. We examined the accuracy of segmentation using 1,000, 5,000, 10,000 and 24,500 small training images. As the number of training images increased, both the F-value and the recall improved, whereas the precision remained relatively consistent (Fig. 4A). We next examined the difference in segmentation accuracy when different areas of the embryo were used for training images (Figs. 4B-D). Cells in the center of the image were of a different type to those in periphery in relation to clarity, area, and oblateness. Thus, we measured the accuracy of segmentation when an image from each region was used for training. We found that F-values were high when the same type of images were used for training, and recall and precision were high when periphery and center images were used, respectively. These results suggest that accuracy is improved when similar images are used for training, and that periphery images guide over-segmentation, while center images guide under-segmentation.

**Fig. 4.**
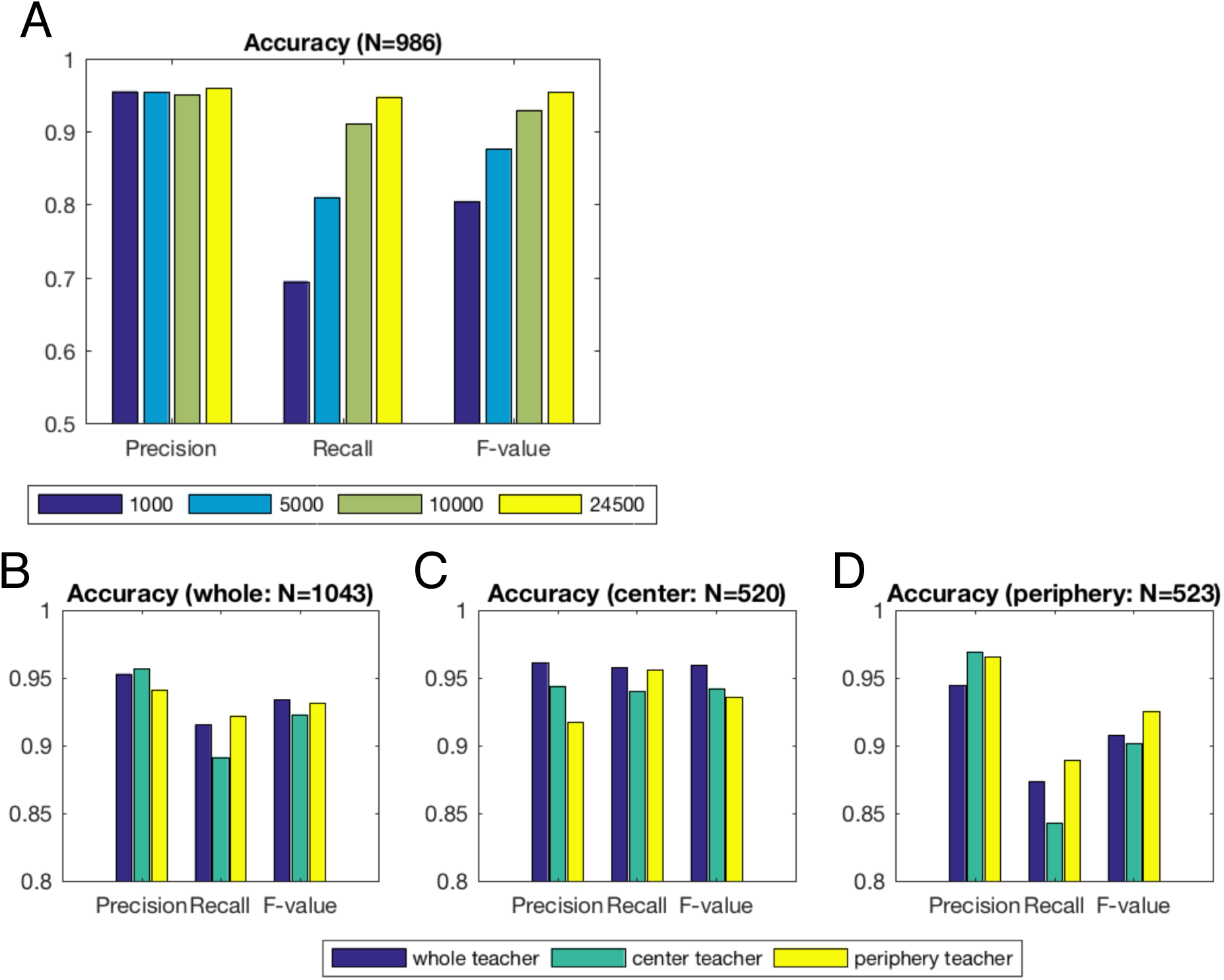
Comparison of precision, recall, and F-value in segmentation under changing conditions. (A) Precision, recall, and F-value with CSML segmentation in relation to the number of training images used. Increased numbers of training images improved both F-value and recall, but precision was largely unchanged. (B-D) Precision, recall, and F-value with CSML segmentation when using the whole region (B), center region (C) or peripheral region (D) of an embryo as training images. Cells in center of the image differ from those in periphery in their clarity, area, and oblateness. Segmentation was accurate for the whole segmentation image, center of the image, and periphery of the image when training was performed using the same type of images (whole image, center of the image, and periphery, respectively). In all trainings, we used 10,000 training images.

For the whole image, the F-value was better when using the whole teacher image, used in training step, rather than using partial images (Fig. 4B). Recall was better when using the whole teacher image rather than using the center, but worse than using the periphery. Precision was better when using the whole teacher image rather than using the periphery, but worse than using the center. In the center of the image, the F-value was better when using the center teacher image than when using the periphery teacher but worse than when using the whole (Fig. 4C). Recall was worse when using the center teacher than it was using the periphery or center. Precision was better when using the center teacher than when using the periphery teacher, and worse than when using the whole teacher. In the periphery of an image, the F-value was better when using the periphery teacher (Fig. 4D). Recall was best when using the periphery and was second best when using whole images. Precision was better when using the periphery than when using the whole teacher, and worse when using the whole teacher.

### Generality of advantage in CSML segmentation

In order to validate the application to other cells, we computed the accuracy of segmentation in three independent images. We examined the accuracy of these images using the same image as training (Figs. 5A-C). Images #1 and #2 were obtained from the same embryos used for validation, with the developmental stage of #1 earlier than that of #2. Image #3 was obtained from an independent embryo. The F-value of CSML was the best in all images examined. This suggests that CSML can be used in wide range of cell images; however, it should be noted that segmentation was more accurate when using a teacher image of a similar stage.

**Fig. 5.**
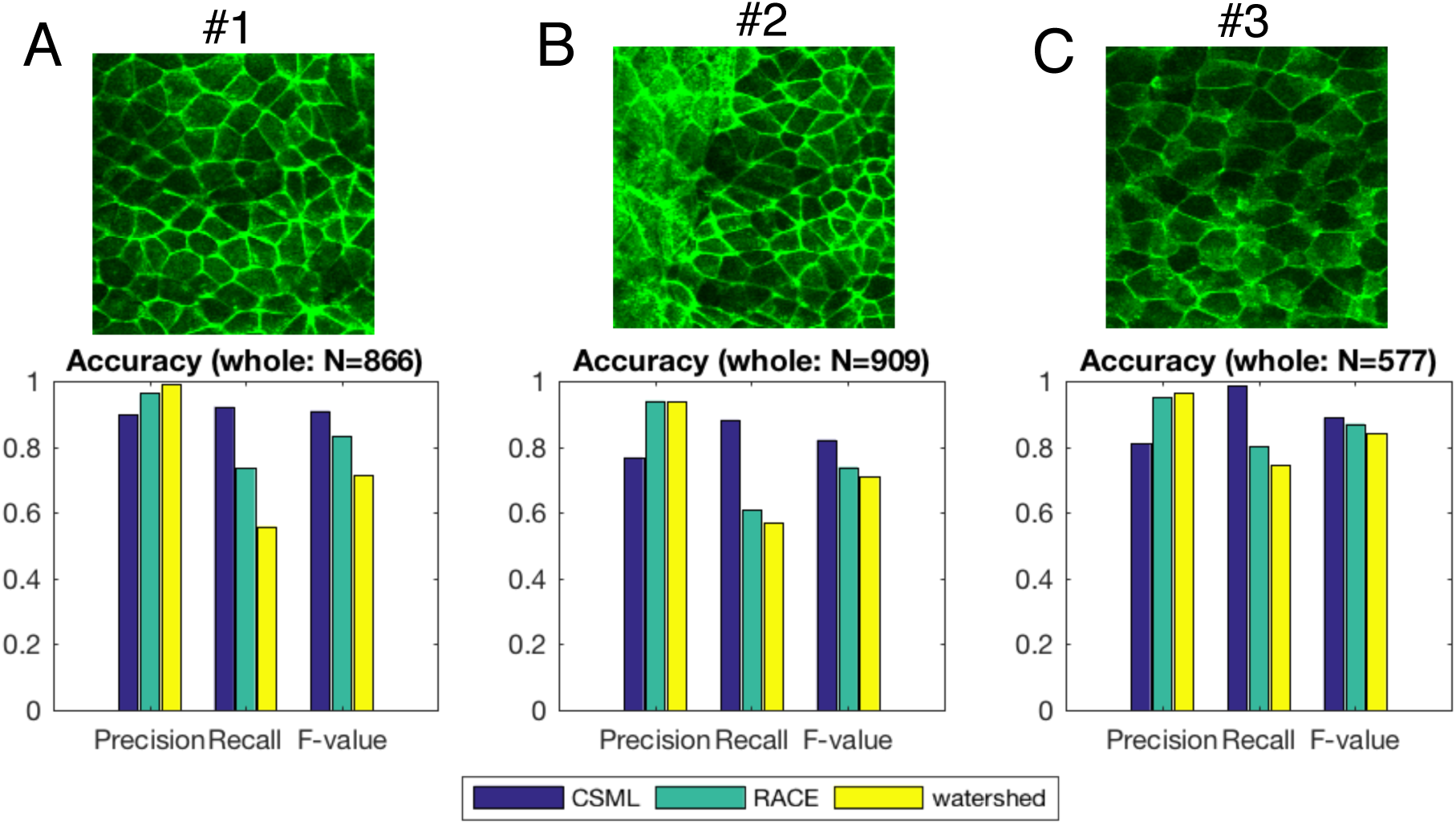
Comparison of precision, recall, and F-value of three independent segmentation images by RACE, watershed, and CSML using the same training images. The F-value of CSML was the best in all of images examined. Segmentation was most accurate in the images most similar to training images.

## Discussion

In this study we established a novel method for automatic cell segmentation using machine learning, and we applied this method to the description of vertebrate embryos. Machine learning is a data-driven technique that avoids the effort of extracting features to construct an algorithm and that provides high accuracy. This is because the FCN classifier can identify membrane regions that previous programs could not, such as when part of a membrane edge is not fluorescent. The main feature of the FCN classifier is its unpooling and deconvolution operations (Noh et al., 2015). In particular, deconvolution may be critical for increased accuracy of segmentation. Deconvolution outputs an enlarged and dense activation map (Noh et al., 2015), which avoids separated membrane regions.

The precision of CSML was slightly lower than RACE and watershed, but the recall was vastly higher. This made the F-value of CSML considerably higher than the others. The reason that the precision of CSML was the poorest may be due to false-negative error rather than false-positive error, which was generally common in watershed and RACE, leading to under-segmentation. While CSML demonstrated the closest correspondence to Ground Truth, with the graph of watershed being obviously different, CSML was still not that close in orientation. The effect may be because orientation would be greatly affected if even a small number of cell edges are not recognized.

CSML using a whole image for training showed the best F-value for the whole image, likely due to whole images containing a wide range of information. On the other hand, recall was best when the periphery image was used, where most cells are small and CSML may fail to recognize cells. Precision was best when using center images, where cells are large. As such, CSML tends to not over-segment cells. In the periphery, the F-value was best when using periphery images for training, as these are then similar to the inferred images. Recall and precision were generally the same. In the center, F-value, recall and precision were best when using the whole image. This may be due to cells in the center of training image being much clearer than the one to be inferred in this experiment. This indicates that using whole images for training was better than using center images, even though both sizes were similar. We also evaluated the accuracy of segmentation with several other fluorescent images, and we found that a better F-value was achieved when the training image was of a similar developmental stage.

## Conclusion

In this study we suggest a novel method for cell segmentation using machine learning. This application enables the segmentation of *Xenopus* embryos, which to date have been difficult to segment with non-machine-learning-based applications. Our application takes 20 seconds per image, which is fast enough for practical use.

To further improve accuracy, regression might be used instead of classification to detect the membrane regions as a connected edge. If large values are set in the center of membrane edges and small values in the periphery of membrane edges, the classifier can be preferentially optimized to detect the center line of edges, which promotes more connectivity amongst inferred edges.

CSML is open-source, user-friendly software. Parameters are changeable, and CSML is available for Windows and MacOS. The program is distributed on https://github.com/RickyOta/CSML, and CSML is on https://github.com/RickyOta/CSML/releases.

## Acknowledgement

We thank Ryo Nakabayashi for help creating the CSML software. This study was supported by KAKENHI (Grant No: 26440115) to TM.

